# Outer epidermal edges mediate cell-cell adhesion for tissue integrity in plants

**DOI:** 10.1101/2025.04.03.646819

**Authors:** Özer Erguvan, Adrien Heymans, Asal Atakhani, Elsa Gascon, Richard S. Smith, Olivier Ali, Stéphane Verger

## Abstract

Cell-cell adhesion in plants is generally thought to be primarily mediated by the middle lamella, a supposedly thin adhesive layer of the cell wall. Here, we challenge this view. Through computational simulation we found that outer edges of cellular interfaces of the epidermis are hotspot for cell separating tensile stress. Characterization of the ultrastructure of those edges *in planta* revealed that they are locally thickened regions that harbor cellulose lamellae and pectin-based structural continuity across adjacent cells. We confirmed their dominant role for adhesion by studying mutants where those edges are defective and through direct mechanical testing. This reveals the key role of outer epidermal cell edges, rather than the middle lamella, in mediating cell-cell adhesion for tissue integrity in plants.

## Main Text

The mechanisms by which cells physically adhere to each other are well studied in animals, where transmembrane proteins called cell-adhesion molecules form adhering clusters like adherens junctions or focal adhesions (*1*, *2*). However, adhesion is fundamentally different in plants, where cells are surrounded by a thick cell wall (*3–7*). The primary plant cell wall is mainly composed of hemicelluloses, pectins and structural proteins that are secreted to form an amorphous matrix (*8*, *9*) and cellulose that is synthesized at the plasma membrane in a rather organized manner (*10*), forming a layered cross lamellate structure (*11*, *12*). Cell-cell adhesion in plants is generally depicted as individual cell surrounded by their cell wall, adhering to each other through a thin pectin-rich layer called the middle lamella (ML) (*5*, *13–15*). This commonly accepted view considers the ML as the sole or primary structure holding plant cells together.

The ML’s importance has been supported by the fact that pectin degradation or synthesis deficiency generally leads to loss of cell-cell adhesion (*4*). Nevertheless, this does not hold in all cases. For instance, the adhesion-defective phenotype of the pectin-deficient mutants *quasimodo1* and *2* is almost fully rescued by a suppressor mutation in *ESMERALDA1*, without restoring pectin levels (*16*). Interestingly, subsequent work has shown that the *quasimodo* mutants also show cellulose synthesis and cuticle deficiency (*17*, *18*). Similarly, cell separation events generally require a combination of enzymes acting not only on pectin but also cellulose and hemicelluloses (*19–21*). Furthermore, adhesion in plants is established during cell division (*6*). A cell plate forms inside the cell, fusing with the mother cell’s wall. After division, daughter cells share a common cross wall that is likely the basis of the future ML. Then each cell can resume cell wall deposition within their new boundaries and create their individual walls (*22*– *24*). Since adhesion at the ML is established during cell division rather than by *de novo* cell contact, it can be argued that the ML does not actually “stick” cells together in the same way as cell-adhesion molecules do for animal cells. In turn, we may ask what parts of intercellular junctions are key for holding cells together.

In this reasoning, it is crucial to consider the forces at play. Adhesion is ultimately a question of mechanics, where adhesion strength counterbalances separating tensions (Fig 1 A) to maintain tissue integrity (*3*). Geometry and topology play an important but complex role in determining the local distribution of stress and strain at different scales (*25–28*). In turn, we can question whether all tissues, cell layers, cell faces or even cell wall subdomains need to display strong adhesion, or if it is sufficient to reinforce specific hotspots to ensure an organism’s multicellular integrity.

**Fig 1.**
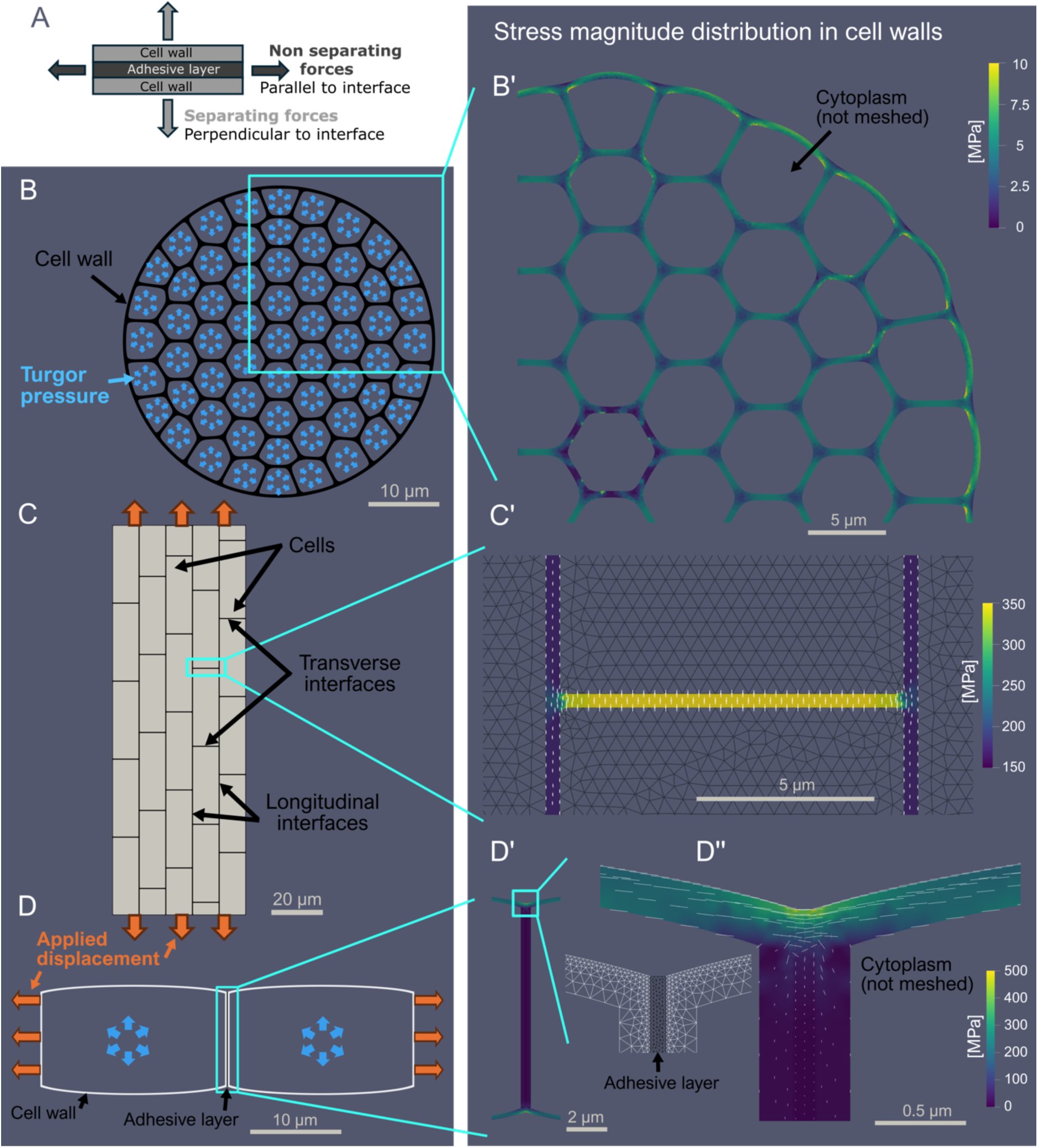
Finite Element Method (FEM) analysis points to the outer epidermal edges of transverse cellular interfaces as hotspots for cell separating tensile stress. (A) Conceptual representation of the “Adhesive layer” and “separating forces” concepts where forces acting perpendicular to the layer could in principles lead to separation at the adhesive layer interface (separating forces), while forces acting parallel would primarily lead to extension (non-separating forces). (B) 2D mesh representing the cross section through a group of cells. (C) 2D mesh representing the surface of an epidermis with staggered cell files, segmented into two subdomains: the cells and the “adhesive layer” (interfaces) between cells. (D) 2D mesh of a longitudinal section representing two epidermal cells with two subdomains: the cell wall and the “adhesive layer” at the interface. In (C) and (D), the Young modulus of the “adhesive layer domain is half the value of the cell or cell wall domain. In (C) and (D), Dirichlet boundary conditions are used on the short sides of the mesh to apply longitudinal stretching (Orange arrows) and reproduce tissue scale directional tension. In (B) and (D), Neumann boundary conditions are used in the cells to apply outward pressure (Blue arrows) and reproduce the effect of turgor pressure. (B’) Heatmap showing stress magnitude distribution over a close-up of the upper right section of the mesh from (B) after applying the boundary conditions. (C’) Heatmap showing stress magnitude distribution over a close-up of a transverse interface from the epidermal surface mesh from (C) after applying the boundary conditions with the two major components of the stress tensors showing the anisotropy of the stress. Note that while the simulation is run on the full meshed area, for illustration purposes, here we only display stress distribution in the adhesive layer (see full simulation in Fig S1). (D’) Heatmap of stress magnitude over a close-up of the anticlinal cell wall from the longitudinal section mesh form (D), while (D’’) shows a close-up of the edge region of the cell junction with the two major components of the stress tensors showing the anisotropy of the stress. On the side, the mesh illustrates the subdomain discretization in the edge region. See corresponding full mesh and strain heat map along with additional supporting simulation output in Fig S1.

### Outer epidermal edges of transverse cellular interfaces are hotspots for cell separating tensile stress

To investigate if there are such strategic adhesion reinforcements points for plant tissues, we first turned to computational simulations using the Finite Element Method (FEM). We simulated the impact of turgor pressure and/or tissue level stretching on the distribution of stress and strain in three simple 2D models with scopes ranging from the organ to the cell wall scale (Fig 1 B-D and S1). At the organ scale, with a mesh conceptually representing a group of fully adhering turgid cells (Fig 1 B), we observed that the stress level was overall the highest in the outermost cell wall layer (Fig 1 B′ and S1 C and D), as previously proposed (*29–31*). This suggests that adhesion may need to be specifically reinforced at the outer epidermal surface. We next investigated the role of anisotropic tensions on such surface layer at the tissue scale. Plant tissues are often exposed to strongly anisotropic stress patterns (*32*, *33*) which may differentially affect stress distribution at cell interfaces which are parallel or perpendicular to the stress. We thus simulated an outer epidermal surface layer as a 2D mesh made of staggered solid cells glued together by a softer “adhesive layer” (Fig 1 C) and applied longitudinal stretching (Fig S1 E and F). Focusing on the stress distribution in the adhesive interfaces between the cells, revealed that transverse interfaces (relative to the stretching axis; Fig 1 C) were exposed to high separating stress (stress that is perpendicular to the interface; Fig 1 A), while longitudinal interfaces were exposed to less stress and in a direction that does not threaten to separate cells (Fig 1 C′ and S1 E’-F’). Finally, at the cell wall level, we simulated the stretching of two pressurized cells adhering through an ML-like layer to study stress distribution at their ML-like interface (Fig 1 D). We observed a very clear pattern where the exposed edges were under high separating tensile stress (Fig 1 D′′), while most of the inner intercellular interface (equivalent to the bulk of the ML as opposed to the epidermal surface studied in the second model) appeared to be under almost no stress (Fig 1 D′ and S1 I-M). Overall, our models suggest that stress globally propagates to the outer epidermal surface (Fig 1 B′), that directional tensions threaten to separate transverse junctions (Fig 1 C′), and that edges of adhesive interfaces are stress hotspots rather than the bulk of the ML (Fig 1 D′′). Taken together, these simulations strongly suggest that the outer epidermal edges of transverse intercellular junctions (relative to stress axis) are hotspots for cell separating tensile stress. In turn, such hotspots may harbor specific structures aimed at dissipating the stress and/or reinforcing cell adhesion for supracellular tissue integrity maintenance.

### Ultrastructural characterization of the epidermal surface reveals supracellular material continuity across intercellular gaps

To identify putative adhesion reinforcing structure, we used *Arabidopsis thaliana* dark-grown hypocotyl as a model since its very highly anisotropic growth has been proposed to lead to high longitudinal tensile stress in the epidermis ((*32*, *33*); Fig 2 A and B). In the adhesion defective mutants *quasimodo2-1 (qua2-1*) and *actin related protein 2-1* (*arp2-1*), that show respectively strong and mild adhesion defects, this leads to the formation of gaps between epidermal cells as revealed using confocal microscopy and propidium iodide (PI) staining of the cell wall (Fig 2 C-G). Those gaps generally form at anticlinal transverse intercellular junctions, perpendicular to the growth axis and growth-derived tensile stress (*34*). Interestingly, using cryo-Scanning Electron Microscopy (cryo-SEM) on *qua2-1* and *arp2-1* mutants, we can observe what appears to be a partially ruptured supracellular layer of material that spans across adjacent cells when there are mild cell separations (Fig 2 H-J). To further confirm these observations, we used a correlative confocal and Atomic Force Microscopy (AFM) approach. This led to similar observations, where the gaps visible by confocal microscopy were partially covered by a material layer made obvious with AFM topological mapping of the surface (Fig 2 K-N and S2). Interestingly, the PI staining used for confocal microscopy also revealed signal continuity across separated cells as shown in orthogonal views of the 3D stacks (Fig S2). Overall, these observations uncover the presence of supracellular material continuity at the periclinal outer surface, which is made visible across partially of fully separated adjacent cells.

**Figure 2:**
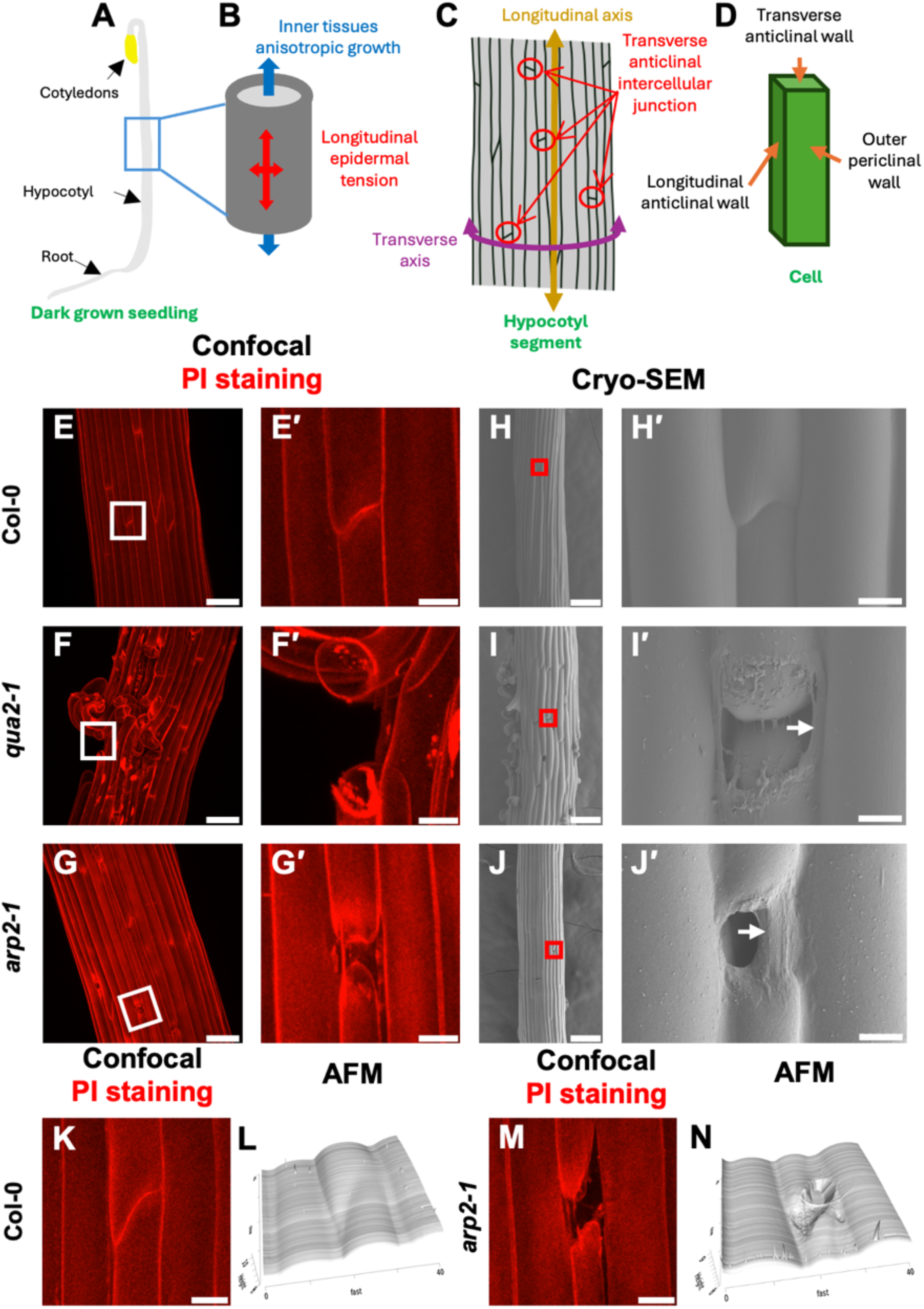
Confocal, Cryo-SEM and Correlative Confocal-AFM reveal a supracellular outer epidermal cell wall layer. (A) Schematic representation of a dark-grown *Arabidopsis thaliana* seedling. (B) Schematic representation of growth and tensions patterns on a dark-grown hypocotyl segment. (C) Schematic representation hypocotyls longitudinal and transverse axes and highlighting the presence of transverse anticlinal intercellular junctions on a cellularized dark-grown hypocotyl segment. (D) Schematic representation the outer periclinal wall and longitudinal and transverse anticlinal walls on a cell. (E-G and E’-G’) Representative Z-projection (maximal intensity) images of confocal stacks from 4-day-old dark-grown hypocotyls stained with propidium iodide (note: white square in E-H indicates where the images are cropped from for E’,H’) for Col-0 (E, E’), *qua2-1* (F, F’) and *arp2-1* (G, G’). (H-J and H’-J’’) Representative Cryo-Scanning Electron Microscopy (cryo-SEM) micrographs from 4-day-old dark-grown hypocotyls for Col-0 (H,H’), *qua2-1* (I,I’) and *arp2-1* (J,J’). Red square in H-J indicate where the high-resolution cryo-SEM images (H’-J’) were acquired and white arrow in I’ and J’ indicate the supracellular outer epidermal cell wall layer. (K and M) Representative Z-projection (maximal intensity) images of confocal stacks from 4-day-old dark-grown hypocotyls stained with propidium iodide showing epidermal cell wall junctions for Col-0 (K), *arp2-1* (M) and (L and N) corresponding Atomic Force Microscopy (AFM) high-resolution 3D surface topology images showing epidermal cell wall junctions for Col-0 (L), *arp2-1* (N). Note that these observations result from a correlative microscopy approach and display the same junction (K/L and M/N with the two modalities. Scale bars: (E-G) 50µm, (E’-G’) 10µm, (H-J) 100µm, (H’-J’) 5µm, (K and M) 10µm.

### Increased local thickness at outer epidermal cell edges can dampen cell separating stress

Next, we turned to Transmission Electron Microscopy (TEM) to characterize the ultrastructure and components of those transverse intercellular junctions. We generated longitudinal sections on Col-0 and *arp2-1* (Fig 3 A-C, G-I). We observed that the cross-section of the transverse outer epidermal edge (OEE; See Fig 3 C, D, G, I) roughly resembles a pseudo-triangle or deltoid curve (triangle with concave sides), showing that the OEE is a structure that is locally thickened compared to the rest of the outer epidermal wall. We quantified local cell wall thickness (Fig 3 H, Table S1) on Col-0 junctions and found that the edge (OEE) is on average

**Figure 3:**
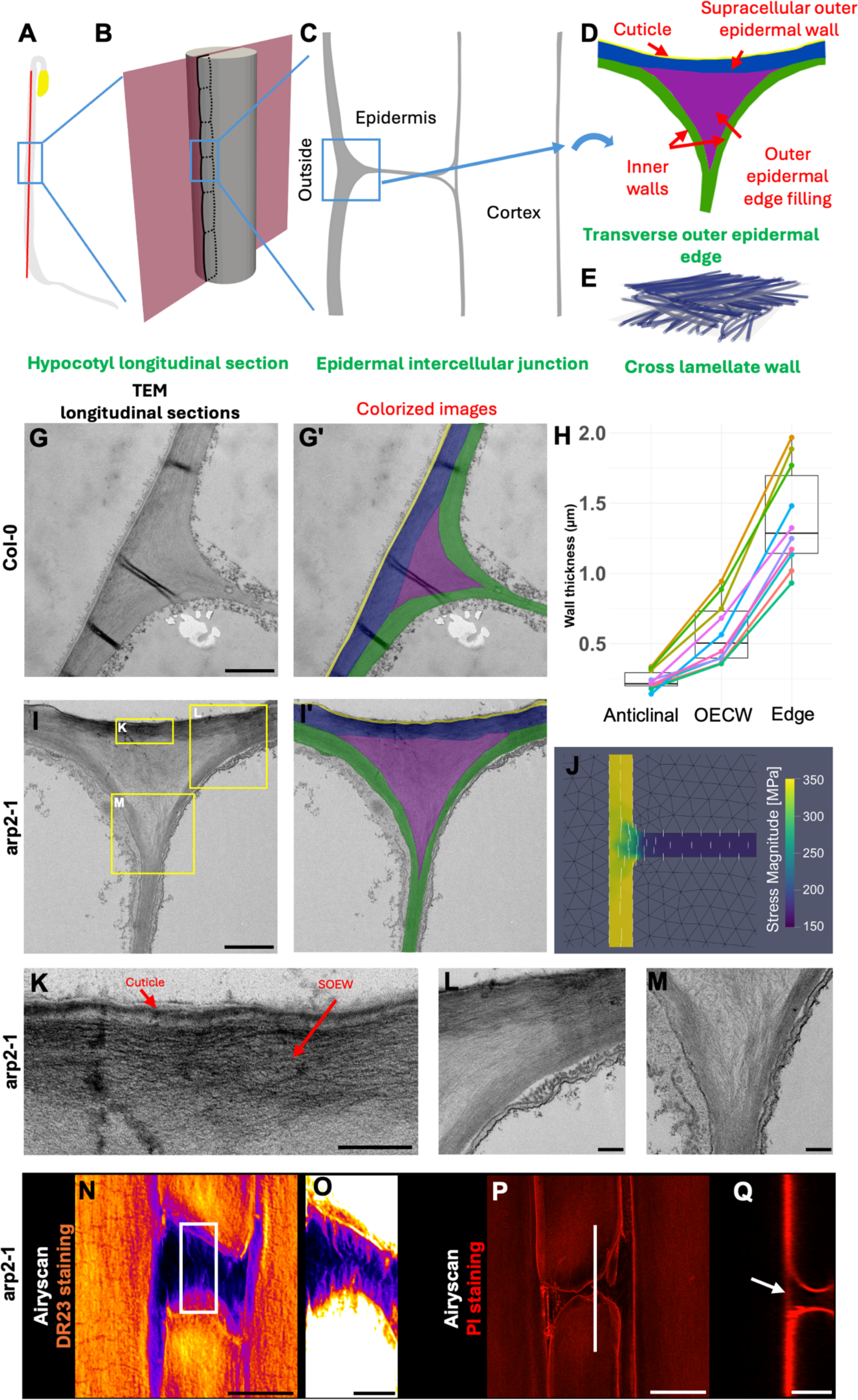
TEM reveals increased local thickness and cellulose- and pectin-based supracellular outer epidermal wall continuity across transverse junctions. (A) Schematic representation of a dark-grown *Arabidopsis thaliana* seedling with a red line representing the axis of sample sectioning for Transmission Electron Microscopy (TEM) preparation. (B) Close up schematic representation of a dark-grown hypocotyl segment and the plan of sample sectioning with epidermal cell outlines for a cell file along the sectioning plan. (C) Schematic representation of a thin longitudinal section through epidermal and cortical cells around an intercellular junction on the epidermal cell layer of a dark-grown hypocotyl. (D) Schematic representation of a transverse outer epidermal edge, annotated subdomains as identified in G’ and I’). (E) Schematic representation of cellulose microfibrils cross lamellate organization. (G-I) Representative TEM micrographs of longitudinally sectioned transverse outer epidermal edge from a (G) 2-day-old Col-0 and (I) 4-day-old *arp2-1,* dark-grown hypocotyls. (G’ and I’) are colorized version of (G and I) highlighting the different subdomains of the cell wall identified in our observations: yellow: cuticle, blue: supracellular outer epidermal cell wall (“SOEW”), pink: outer epidermal cell edge filling (“OEEF”) and green: inner walls (“IWs”) and as described in (D). (H) Box plot of the cell wall thickness quantification from 10 transverse outer epidermal edge regions of 4-day-old dark-grown hypocotyls. Points colors and the lines linking them correspond to data measured on the same edge region, highlighting the consistent trend of differences in cell wall thickness. (J) Heatmap showing stress magnitude distribution over a close-up of a transverse interface from the epidermal surface mesh and after applying the boundary conditions presented in Fig 1 D and where the Young modulus of the “adhesive” layer domain in this case is twice the value of the cell domains. The two major components of the stress tensors are plotted with segments to show the anisotropy of the stress. Note that while the simulation is run on the full meshed area, for illustration purposes, here we only display stress distribution in the adhesive layer (see full simulation in Fig S1 G-H). (K-M) Close ups from I (See yellow rectangles) highlighting the cross lamellate and organized ultrastructure of the SOEW along with the cuticle (K), the SOEW/IW periclinal interface (L) and the IW/IW anticlinal interface (M). (N and Q) Representative Z-projection (maximal intensity) Airyscan confocal images from 4-day-old dark-grown *arp2-1* hypocotyl stained with Direct Red23. Note that we applied a “fire” look up table to highlight the presence of fibrillar structures across the partially separated cell junction and that in the close up (O) we have increased the contrasts to further highlight those structures present across separated cells. (P) Representative Z-projection (maximal intensity) Airyscan Confocal images from 4-day-old dark-grown *arp2-1* hypocotyl stained with Propidium Iodide. (Q) Orthogonal view through the 3D confocal stack (See white line in (P)), highlighting the presence of a continuous PI-stained outer epidermal wall layer across mildly separated cells. Scale bars: (G-I) 5µm, (K-M) 200nm and (N-Q) 10µm.

2.6 times thicker than the adjacent outer epidermal walls. Thickening of the cell wall at these predicted stress hotspots might protect the junctions from separating tensile stress. To explore this idea, we used our simulation of the epidermal surface presented in Figure 1 C, doubling the theoretical thickness of the interfaces compared to the cell domains. In turn, compared to the original simulation, we observed a rather opposite pattern in terms of stress intensity at the interfaces, while the main orientation of the stress remained the same (Fig 3 J). This suggests that thickened intercellular edges can indeed buffer the stress that would threaten to separate transverse interfaces. Instead, thickened edges divert the stress to longitudinal interfaces where the stress direction does not threaten adhesion (Fig3 J and S1 G-G’).

### Cellulose- and pectin-based supracellular outer epidermal wall continuity is present across transverse junctions

Next, we took a closer look at the cell wall ultrastructure of the transverse OEE. In our TEM images, we could distinguish the characteristic cross lamellate structure of the cell wall ((*12*); Fig 3 E, G, I and K). On one hand we observed as expected that those lamellae were visible at the junctions, spanning from the periclinal to anticlinal wall of adjacent cells, where they meet at the putative location of the ML (Fig 3 I and M). On the other hand, we observed cell wall lamellae along with the cuticle proper spanning across the outer epidermal surface of adjacent cells (Fig 3 G, I, K and L), while the center of the pseudo-triangle-shaped OEE appeared less clearly organized (Fig 3 D, G, I and K-M). Those structures were also observed and appeared even more obvious in *arp2-1* samples (Fig 3 I, K-M and 4 F-I). Here we propose to name these structures “inner walls” (IWs), “supracellular outer epidermal wall” (SOEW) and “outer epidermal edge filling” (OEEF) respectively (Fig 3 D). Thus, along with the SEM, AFM and confocal observations (Fig 2 E-N and S2) these TEM images confirm the presence of a continuous layer of material across adjacent cells. A close-up view on a SOEW clearly shows both cuticle and layered fibrous material, indicating the presence of cellulose-based lamellae across neighboring cells (Fig 3 E and K). We further confirmed this observation using high-resolution imaging with Airyscan and joint deconvolution on *arp2-1* with direct red 23 (DR23) and PI staining highlighting the presence cellulose and pectin in the SOEW across mildly separated junctions (Fig 3 N-Q).

**Figure 4:**
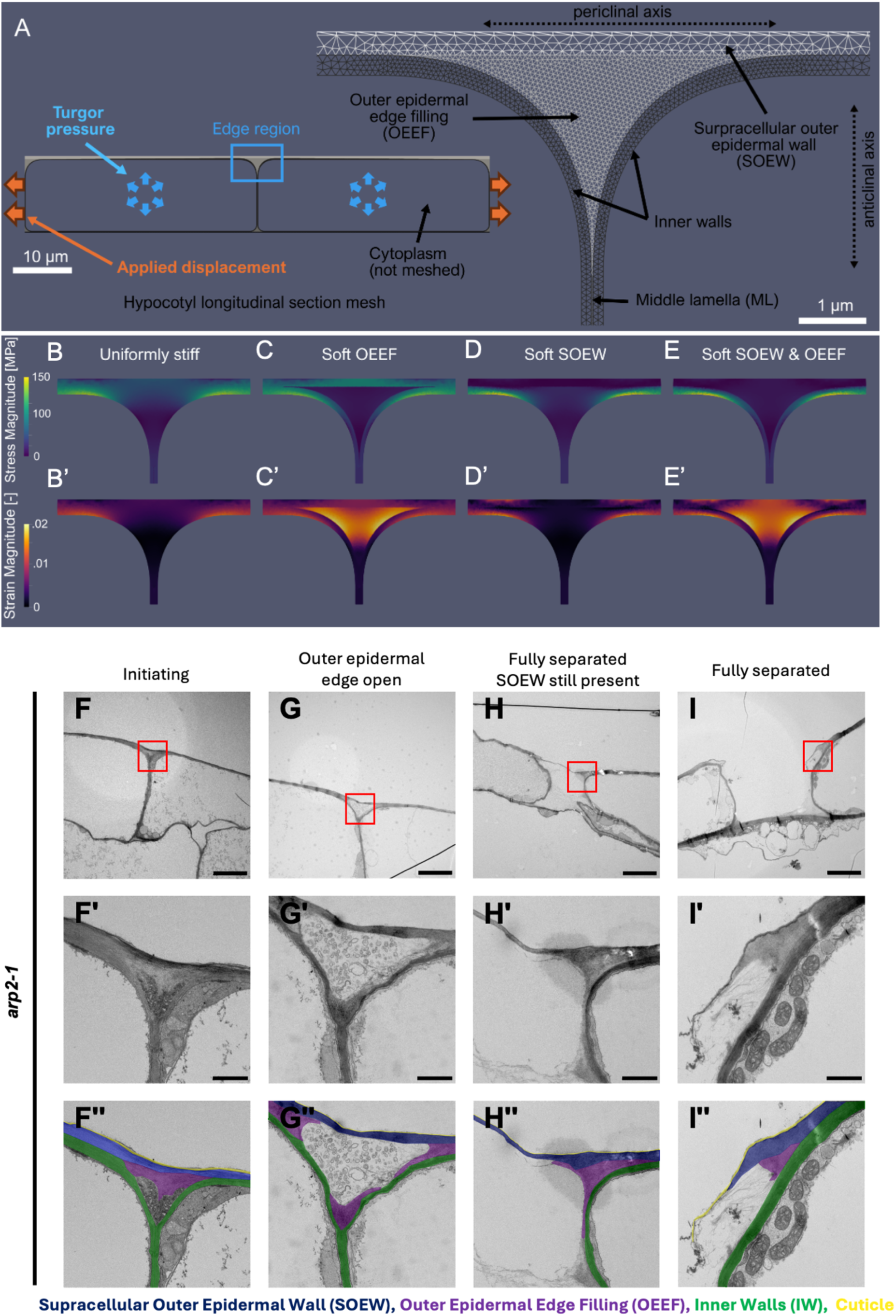
FEM and TEM reveal the process by which cell separation may initiate in the outer epidermal edge filling (OEEF) and propagate to the supracellular outer epidermal wall (SOEW). Finite Element Method (FEM) analysis of stress and strain distribution on a longitudinal section of hypocotyl epidermal cells under varied subdomain Young’s modulus conditions. (A) 2D mesh of the hypocotyl longitudinal section on the left, with a close-up on the right of the edge region at the outer epidermal cell junction based on the structure described in Figure 3. Four subdomains are identified: the Supracellular Outer Epidermal Wall (SOEW), the Outer Epidermal Edge Filling (OEEF), the Inner Walls, and the Middle Lamella (ML). (B– E) Stress and (B’-E’) strain distribution at the edge under both a fixed longitudinal displacement from both ends, stretching the mesh, and turgor pressure. Four simulated scenarios are represented with varied Young’s modulus values. Unless specified otherwise, Young’s modulus for all subdomains is set to 10 MPa, with specific scenarios softening subdomains to 1 MPa. Stress magnitude heatmaps are shown in (B) with uniformly stiff properties, (C) with OEEF softened, (D) with SOEW softened, and (E) with both OEEF and SOEW softened. Strain magnitude heatmaps are shown in (B’) with uniform properties, (C’) with OEEF softened, (D’) with SOEW softened, and (E’) with both OEEF and SOEW softened. See additional supporting simulation output in Fig S3. (F-I) Representative low magnification (2000X) Transmission Electron Microscopy (TEM) micrographs of longitudinally sectioned 4-day-old dark-grown *arp2-1* hypocotyls showing different stages of cell-cell adhesion defects: (F) Initiating, (G) Outer epidermal edge open, (H) Fully separated while Supracellular Outer Epidermal Wall (SOEW) is still present, (I) Fully separated. (F’-I’) Representative high-resolution (11000X) TEM micrograph from the same samples as (F-I) where red squares show where the high-resolution images were acquired. (F’’-I’’) are colorized version of (F’-I’) highlighting the different subdomains of the cell wall identified in our observations: yellow: cuticle, blue: supracellular outer epidermal cell wall (“SOEW”), pink: outer epidermal cell edge filling (“OEEF”) and green: inner walls (“IWs”). Scale bars: (F-I) 5µm, (F’-I’) 1 µm.

While the IWs and the ML are the cell wall structures classically depicted in plant cell adhesion conceptual models (*5*, *13–15*), our observations now reveal additional structures that are strategically placed with regard to the cell separating stress hotspots that we identified through computational simulation (Fig 1 and S1). Technically, the OEEF is likely a continuation of the ML, yet the ML being a sheet in 3D, while the OEEF forms a wedge at the outer edge of the cellular interface, is a major geometrical difference that has mechanical implications, making them distinct “domains”. Importantly, we unveil the presence of a supracellular cell wall layer (the SOEW), likely derived from cell division (see introduction and discussion) leaving maternal wall continuity spanning across daughter cells.

### The SOEW and OEEF can shield the middle lamella from separating tensile stress

We next investigated how such cell wall subdomains may in theory contribute to adhesion maintenance. We created a mesh with realistic dimensions and subdomains capturing the complexity observed (Fig 4 A and Table S1). The full mesh consists of two turgid and longitudinally stretched cells where the outer epidermal edge is made of 4 subdomains (IWs, ML, SOEW and OEEF) as represented in Fig 4 A.

On a uniform mesh, where all domains are uniformly stiff, the stress was globally higher in the periclinal wall region, but locally lower just above and at the anticlinal wall (Fig 4 B and S3 B and D). This confirms our previous simulations that by geometry only the OEE can shield the ML from separating forces (Fig 3 J). It also confirms that the ML domain as we define it, should experience negligible separating stress and may in turn only have a minor mechanical role for maintaining adhesion. We next questioned the relative contribution of the subdomains for the stress and strain distribution at the outer epidermal edge. We simulated softened domains by decreasing by ten times their stiffness as a proxy for weaker adhesion strength. The rational being that softer walls relax and redistribute stress to stiffer wall domains as if the softer domain was tearing apart and we interpret the resulting strain as the extent of separation. Softening the ML domain alone did not significantly affect stress distribution (Fig S3 B, C and E). Softening of the OEEF alone revealed large separating strain in the OEEF (Fig 4 C′) while the SOEW became load bearing, experiencing very high stress just above the junction (Fig 4 C). Softening the SOEW alone, consequently decreased the stress in the SOEW at the junction (Fig 4 D) but otherwise had only minor effects on the global stress and strain pattern of the edge (Fig 4 D, D′). Finally, the strongest effect was achieved when both SOEW and OEEF were softened simultaneously (Fig 4 E and E′). The separating strain was consequently strongly increased in the SOEW and OEEF (Fig 4 E′), resembling the situation where the OEEF alone was softened, except that the SOEW was not load bearing anymore (Fig 4 E). In turn separating strain strongly focuses just above the cell junction on the SOEW (Fig 4 E and S3 B’). Thus, on one hand, our simulations suggest that the OEEF is a key structure where cell separating strain or cracks may initiate and propagate if softened. On the other hand, redistribution of the stress to the SOEW when the OEEF is softened further points to the SOEW as a potentially key load bearing structure across adjacent cells.

### Intercellular edges ultrastructure of separating cells reveals the process by which cell separation may initiate and propagate

In our TEM observations on the *arp2-1* mutant, we acquired a range of adhesion-defective intercellular junctions ranging from intact to fully separated. In some of the mildest cases, the OEE appeared generally intact, but we observed a much clearer distinction between the SOEW, IWs and a more loosely packed OEEF (Fig 3 I). Next, we observed cases where what appeared to be a nucleation point of cell wall disruption or fracture had formed at the base of the OEEF (Fig 4 F). In the third example we saw what appears as a large gap instead of the OEEF (Fig 4 G). The structure strikingly resembles intercellular spaces found within inner tissues. In the next example, the cells appeared fully separated at their anticlinal face while we again observed a continuous SOEW across the cells (Fig 4 H). In the last example, cells were fully separated, the SOEW was ruptured but the IWs appeared still intact (Fig 4 I).

Interestingly, this range of adhesion defective junctions reveals the process by which cell separation may initiate and propagate in those transverse edges and strikingly fit our model predictions. Fracture nucleation may happen at the base of the OEEF when it is weakened in the adhesion defective mutant background. It could then propagate to the full OEEF followed by the middle lamella, until cells fully separate and the SOEW may rupture last.

### Combined impairment of the SOEW and OEEF correlates with enhanced adhesion deficiency

The *qua2-1* mutant was previously shown to not only be deficient in pectin synthesis but also cellulose and cuticle due to secondary responses (*17*, *18*). On the other hand, the suppressor line *qua2-1 esmd1-1* shows a strongly rescued adhesion phenotype while still having similar pectin deficiency as *qua2-1* (*16*). We hypothesized that in *qua2-1* the ML, OEEF and SOEW may be compromised, while in *arp2-1* and *qua2-1 esmd1-1* only the ML and OEEF would be.

To determine this, we used TEM to examine the OEE of transverse junction on 2-day-old dark-grown seedlings, before the growth-derived epidermal tensions pull cells apart. Interestingly, at such early stage, Col-0, *arp2-1*, *qua2-1 esmd1-1* junctions appeared fully intact and structurally undistinguishable from each other (Fig 5 A, B and D). On the other hand, in *qua2-1* the junction structural organization appeared already noticeably altered, with a general collapsed and irregular appearance (Fig 5 C). Next, we used DR23 staining and airyscan confocal microscopy on 4-day-old dark-grown seedlings and looked at partially separated cell junctions. One clear difference was that, while for *arp2-1* and *qua2-1 esmd1-1* we could generally observe fibrillar structures across mildly separated cells (Fig 5 F and H), in *qua2-1* this cellulose staining generally revealed rather globular structures (Fig 5 G). Overall, these observations suggest that in *qua2-1*, the cellulose network within the SOEW is poorly organized and structurally impaired compared to *qua2-1 esmd1-1* and *arp2-1*. As previously reported the *qua2-1* mutant shows a much stronger adhesion defective phenotype than *qua2-1 esmd1-1* or *arp2-1* mutants (Fig 2 E-G and (*16*, *35*)). These experimental observations fit our model predicting that a combined impairment of the SOEW and OEEF would have a stronger effect on adhesion than impairment of OEEF/ML alone.

**Figure 5:**
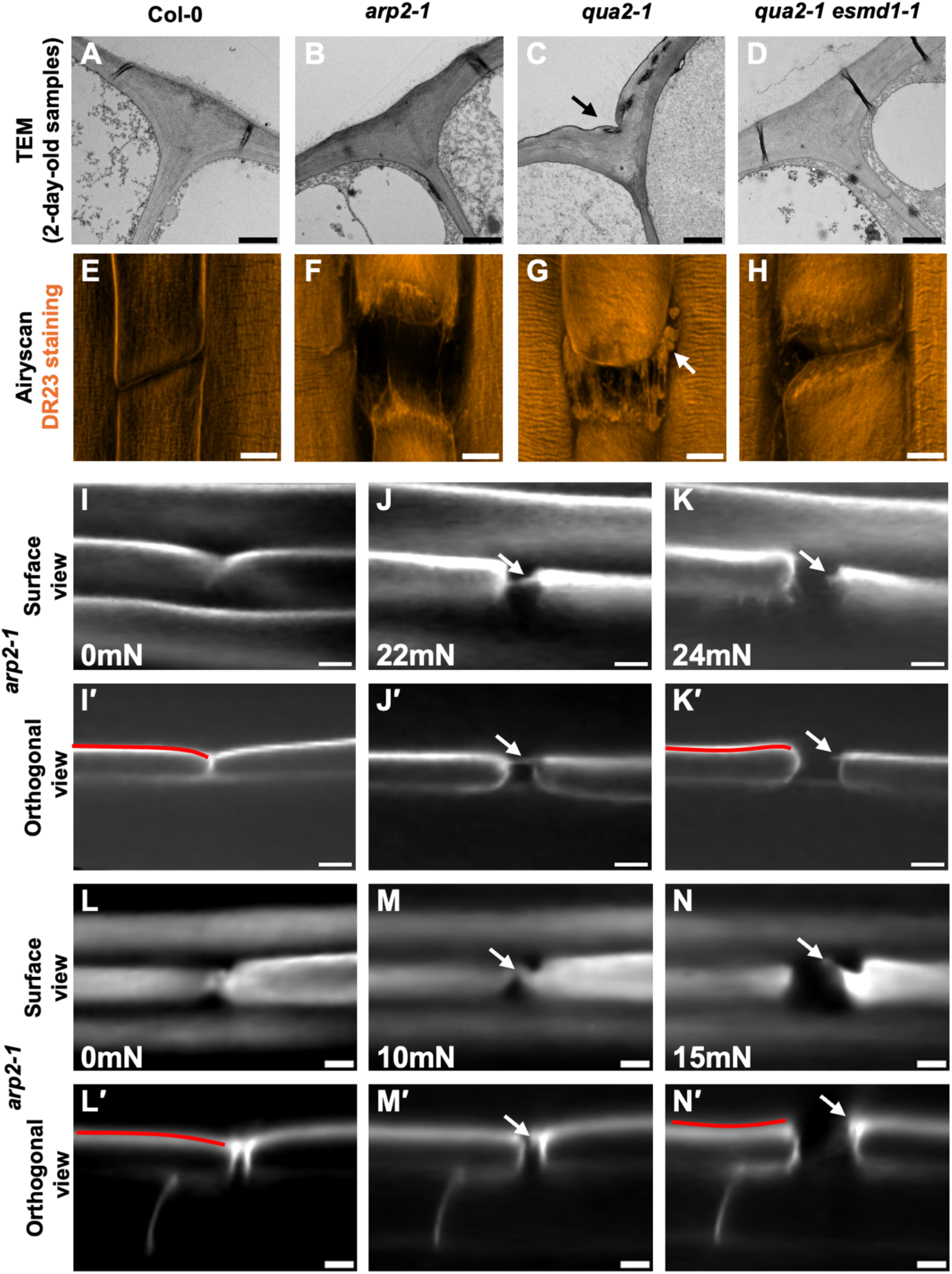
The SOEW and OEEF physical integrity contribute to cell adhesion maintenance in plants. (A-D) Representative Transmission Electron Microscopy (TEM) micrographs of longitudinal-sectioned 2-day-old dark-grown hypocotyls showing outer epidermal cell wall junctions for Col-0 (A), *arp2-1* (B), *qua2-1* (C) and *qua2-1 esmd1-1* (D). (E-H) Representative Z-projection (maximal intensity) Airyscan Confocal images of 4-day-old dark-grown hypocotyls hypocotyl stained with Direct Red23, showing outer epidermal cell wall junctions for Col-0 (E), *arp2-1* (F), *qua2-1* (G), *qua2-1 esmd1-1* (H). (I-N) Representative surface view and (I’-N’) orthogonal view images generated using spinning disk confocal images from Propidium iodide (PI) stained 4-day-old dark grown *arp2-1* hypocotyls that were stretched using an automated micro-extensometer. White arrows indicate the supracellular outer epidermal wall (SOEW). In I and L when 0mN force is applied, J and M when 22mN and 10mN force is applied respectively, K and N when 24mN and 15mN force is applied respectively. Red lines in I’ and L’ shows the convex outer epidermal wall curvature before SOEW ruptures, and in K’ and N’ shows the concave curvature after rupture. See additional data in Video S1. Scale bars: (A-D) 1µm, (E-H) 5µm, (I-N and I’-N’) 10µm

### The SOEW is a load bearing structure spanning across adjacent cells

Finally, we used an automated micro-extensometer coupled with a confocal microscope to mechanically stretch hypocotyls of *arp2-1* and observed how the gaps form between cells during stretching. We used PI staining and acquired 3D stacks to image the cell-cell interface, including the SOEW, and selected junctions where cells were not already visibly separated before the applied stretching. In several cases the stretching did lead to the opening of junctions where we could see the anticlinal cell interface separating while still clearly distinguishing the presence of a SOEW structure still largely or partially intact across the separated cells (Fig5 I-N and movie S1). This fits our TEM observations where the SOEW appears to remain even when cells are largely separated below (Fig 4 G). With increased force, the SOEW also ruptured later-on leading to a sudden opening of the gap and apparent recoil of the outer epidermal surface of the detached cells. This sudden relaxation event is also shown by the change of the curvature of the outer epidermal wall switching from convex before SOEW rupture to concave after rupture (see red lines Fig 5 I’, K’, L’ and N’). Overall, these results experimentally demonstrate that the SOEW can retain significant mechanical strength and play a key mechanical role in maintaining cell adhesion and tissue integrity in plants.

## Conclusion

Here we challenge the widely accepted view that the ML is the main mediator of cell adhesion in plants and instead reveal the presence and key contribution of what we name the Supracellular Outer Epidermal Wall (SOEW) and the Outer Epidermal Edge Filling (OEEF). Those are specifically located where we identified cell separating stress hotspots through computational simulation. We propose that they have a synergistic effect for maintaining adhesion. The thickened structure of the edge plays as stress dissipating role, while the SOEW brings direct load bearing mechanical reinforcement. In turn the OEEF emerges as a key adhesion site, where fracture can initiate and propagate to the ML when weakened.

Pectins can form strong hydrogels (*36*), but in principle cellulose is still orders of magnitude stronger (*11*). Thus, cellulose-based wall domains of the SOEW are likely much stronger than the pectin-rich ML, conceptually forming a strong continuous outer epidermal envelope (*37*). Indeed, the outer epidermal wall in most developing tissues is well known to be significantly thicker than the inner walls (*31*) and inner tissues develop intercellular spaces (*38*) as if cell were not tightly adhering. Note that adhesion reinforcement points are also thought to exist at tricellular junctions and corners of intercellular spaces (*6*), which may regulate intercellular space formation and plant tissue texture (*7*, *39–41*).

A key question that arises is the origin of the SOEW. Cellulose microfibrils are synthesized by cellulose synthase complexes at the plasma membrane (*10*) and thus cannot contribute to deposit cellulose fibers across adjacent cells. Since plant cell adhesion is established during cell division, the SOEW likely derives from the wall of the mother cell, remaining across daughter cells after division (See introduction). This has previously been shown in roots cortex, leaves mesophyll and vascular cambium (*42–44*), but in those cases it appears that the maternal wall is later partially degraded to allow the MLs to join and an air space to form. This could suggest that the SOEW and OEEF on the outer epidermal surface are either not degraded or, are specifically maintained to ensure adhesion and supracellular integrity during growth and development. Interestingly this could be controlled by edge-directed secretion pathway (*27*, *45*, *46*) in response to wall integrity and mechanical feedback form these stress hotspots (Fig 1 and S1 and (*47–51*). In turn, such cell division-derived structure, keeping continuous cell wall structures across adjacent cell is particularly relevant for maintaining adhesion in fast elongating plant tissues that are organized in longitudinal cell files derived from successive anticlinal transverse cell divisions (*52*).

## Acknowledgement

We thank Olivier Hamant and Stéphanie Robert for helpful comments on the manuscript. We thank the facilities and technical assistance of the Umeå Plant Science Centre (UPSC) microscopy facility and the plant growth facility. We also acknowledge the Umeå Center for Electron Microscopy (UCEM) for providing access to the facility and technical support. We especially thank Cheng Choo Lee (Nikki) and Agnieszka Ziolkowska for their invaluable assistance. We thank Simone Bovio for providing AFM training. We thank Sarah Robinson and Mateusz Majda for initial help setting up our micro extensometer. We thank Manuel Petit for proofreading the documentation on the FEM simulations.

## Author contribution

Conceptualization, S.V., O.E. and A.H.

Methodology, O.E., A.H., A.A., O.A., and S.V.

Software, A.H., A.A., R.S.S. and O.A.

Formal Analysis, O.E. and A.H.

Investigation, O.E., A.H. and A.A.

Visualization, O.E., A.H. and S.V.

Resources, E.G. and O.A.

Writing – Original Draft, O.E., A.H, A.A., and S.V.

Writing – Review & Editing, O.E, A.H., A.A., E.G., R.S.S., O.A. and S.V.

Supervision, S.V. and O.A.

Funding Acquisition, S.V.

## Data and material availability

All code and meshes used for the FEM simulations are available in a GitHub repository, executable through Jupyter Notebooks and archived at a Zenodo repository (https://github.com/VergerLab/2D-HypocotylFEM; doi: 10.5281/zenodo.14192237). All the microscopy data used to support this study (including those not displayed in figures) has been deposited at Zenodo on two separate repositories and is publicly available (doi 1: https://zenodo.org/records/15125559; doi 2: https://zenodo.org/records/15124336).

## Supplementary Materials

Figs. S1 to S3 and Tables S1

Movie S1

## Material and Methods

### Plant materials and growth conditions

In this study, *Arabidopsis thaliana* wild-type (Col-0), *qua2-1* (*53*), *arp2-1* (*35*, *54*, *55*) and *qua2-1 esmd1-1* (*16*) were used. The seeds were surface sterilized. Half strength Murashige and Skoog (MS) medium (Duchefa) supplemented with 0.5 g/L 2-(N-morpholino) ethanesulfonic acid (MES) (Sigma-Aldrich) was used as the base medium. For standard growth conditions, 0.8% agar (Duchefa) was used as a gelling agent. For micro-extensometer experiments, 2.5% agarose (VWR) was used as the gelling agent.

Seeds were sown, then first stratified at 4 °C for 2 days in darkness and then light-induced for 4-6 hours in white light (150 μmol/m^2^sec). After that, plates were wrapped with 3 layers of aluminum foil vertically and germinated in an *in-vitro* growth room (16hr light at 22°C/ 8hr dark at 18 °C) for 2 days or 4 days. Seedling age was counted from the beginning of light induction.

### Cell wall staining

For cell contour and pectin staining, propidium iodide (PI, Sigma-Aldrich) was used as described in (*56*). Briefly, plants were immersed in 0.1 mg/ml PI for 6 to 10 minutes in the case of *arp2-1* and Col-0 but not more than 3 minutes in the case of *qua2-1*, and then samples were rinsed with sterile water before mounting with water. For cellulose staining, Direct Red 23 (Sigma-Aldrich) was used as described in (*57*). Briefly, samples were immersed in Direct Red 23 solution dissolved in Phosphate-buffered saline (PBS) with the final concentration 14 mg/ml for 60 minutes under a mild vacuum then kept on a rotator in darkness for 60 minutes. After the incubation, samples were rinsed with PBS before mounting with PBS. Samples were mounted between regular slide and coverslip for classical confocal microscopy, with high performance cover glass (Ziess) for Airyscan, or according to the method described in the confocal-AFM and extensometer experiment sections.

### Confocal microscopy

Laser scanning confocal images were acquired using an upright Zeiss LSM 800 confocal microscope. A long-distance water dipping objective (W Plan-Apochromat 40x/1.0 DIC VIS-IR M27-Water) was used for the correlative confocal-AFM while a water immersion objective (C-Apochromat 40x/1.2 W Korr FCS M27) was used for standard confocal imaging. Z-stacks were acquired using the Zen Blue 3.2 software with the following settings: image size 319.45 x 319.45 µm (1024x1024 pixel), pixel size (XY) 0.31 µm, Z-step size 0.5 µm and without averaging as 16-bit images. Propidium Iodide (PI) excitation was done by using a 561 nm solid-state laser and the emission was acquired in the range of 562-600 nm.

Airyscan confocal images were acquired using an inverted Zeiss LSM 980 confocal microscope equipped with a water immersion objective (D LCI Plan-Apochromat 40X/1.2 Imm autocorr DIC FC). Z-stacks were acquired using Zen 3.8 software as 16-bit images without averaging. To keep the theoretical pixel size of 0.14 µm following settings were used: for Direct Red 23 (Sigma-Aldrich) staining, image size 30.30 µm x 30.30 µm (580 x 580 pixels), Z-step size 0.22 µm, zoom factor 7 and excitation/emission settings of 514 nm and 527-735 nm, respectively. For Propidium Iodide (PI), image size 42.3 µm x 42.3 µm (744 x 744 pixels), Z-step size 0.22 µm, zoom factor 5 and excitation/emission settings of 561 nm and 305-735 nm, respectively. Airyscan Joint deconvolution was applied on all the Z-stacks using the Zen Blue 3.8 software with the following settings: sample structure: standard, maximum iterations 10, quality threshold 0.

Spinning disk confocal images were acquired using an upright Nikon LV100ND CrestOptics Cicero Spinning disk equipped with a long-distance water-dipping objective (Olympus XLUMPlanFL N 20X/1.0 W). Z-stacks were acquired using NIS-Element software with the following settings: 602.94 µm x 602.94 µm (2048 x 2048 pixels), pixel size (XY) 0.29 µm, Z step size 1 µm without averaging as 12-bit. Propidium Iodide (PI) excitation was done using the 561 nm LED and the fluorescence was detected with a filter-based system at 620/60 on a Photometrics prime BSI express camera. Denoise Ai was used on all the acquired confocal Z-stacks using NIS-Element software.

For visualization in figures most of the confocal microscopy 3D data was projected in 2D using a classical maximal intensity projection in ImageJ. Spinning disk microscopy 3D stack used in stretching experiments were further processed in the following way. Stacks were rotated in X, Y and/or Z axis when needed using the TransformJ plugin (*58*)) and cropped in Fiji (*59*)), to center the opening gap and orient it in a way that allows a single slice visualization of its surface as well as XY and XZ orthogonal views through the epidermal cells represented in figure 5 and movie S1.

### Cryo-Scanning Electron Microscopy (cryo-SEM)

The samples were first mounted onto a carbon tape (Ted Pella Inc) on an aluminium stub (Ted Pella Inc). Then the samples were rapidly frozen in liquid nitrogen slush using a cryo preparation unit (Quorum Technology PP3000T). After this, the frozen samples were transferred to a cryo preparation chamber (Quorum Technology PP3000T), and sublimation was done at -90 °C for 10 minutes to remove surface ice crystals. Once sublimation was done, a thin layer of platinum was applied to the samples in the same cryo preparation chamber to prevent beam charging during imaging. The samples were then transferred to the cryogenic field-emission scanning electron microscope (Carl Zeiss Merlin FESEM), with the stage temperature set to -160 °C. All images were acquired with a secondary electron detector, at low beam voltage and current.

### Correlative confocal and Atomic Force Microscopy (AFM)

Samples were first stained with PI as described in the cell wall staining section, then mounted on a glass-bottom petri dish that has grid pattern (35 mm Dish | No. 1.5 Gridded Coverslip | 14 mm Glass Diameter, MatTek) with a thin layer of FLEXBAR Reprorubber® Thin Pour Material (Flexbar Machine Corporation) that does not cover the sample but immobilized the sample. The samples were then immersed in water before imaging with the Zeiss LSM 800 confocal microscope using a long-distance water-dipping objective as described in the confocal microscopy section. At least 2 confocal images were acquired from two different regions of interest along each sample. We then used the brightfield mode and camera mounted on our LSM800 system to acquire a large tile scan area around the sample where both the sample and petri dish grid pattern are visible. After this step, the petri dish was placed on our AFM (see description below) equipped with an upright imaging system. The region of interest was first located using the upright microscope brightfield and camera, guided by the grid pattern on the petri dish. To find the exact cell wall junction that was previously imaged with the confocal microscope, we first acquired low resolution Quantitative Imaging^TM^ (QI) maps at the AFM with the following settings: 120 x 180 µm image size and 60 x 90 pixels (pixel size 2 µm). Once the exact cell wall junction was visible on the low-resolution QI height channel, the high resolution 30 x 30 µm image size, 120 x 120 pixels (pixel size 250 nm) QI data was acquired as described below.

A NanoWizard® 4 XP BioScience AFM (Bruker) equipped with HybridStage^TM^ controlled by JPK SPMControl Software version 7 was used for all AFM experiments. To determine the apparent stiffness of the cantilever used in all the experiments, AFM cantilever calibration was first performed in air using the Sader method (*60*), as described by (*61*)Initially, the contact-free calibration tool in the JPK acquisition software was used to measure the resonance frequency and quality factor. These values were then input into the Sader Method website (https://sadermethod.org) to calculate the cantilever’s spring constant. Once the spring constant was obtained, the method called contact-based in the JPK acquisition software was employed to adjust the cantilever’s sensitivity. With both the spring constant and sensitivity determined, the laser’s sum signal was corrected using the knob to align the signal direction to the photodiode. Finally, the contact-based tool was used again in water to adjust the sensitivity for liquid conditions.

All the AFM experiments were performed in water at room temperature using aluminum-coated TESPA-V2 cantilevers (Bruker), which have a spring constant of 42N/m and a nominal tips radius of 8 nm, with a pyramidal-shaped tip. All the AFM experiments were done using Quantitative Imaging^TM^ (QI) mode with the following settings: setpoint 250nN, pixel time 133 ms, Z length 4.5 µm, speed 79.4 µm/s. 3D topology images were generated from AFM height map and error maps were created by applying Edge Filter function to those height map images in JPK Data Processing Software.

### Transmission Electron Microscopy (TEM)

Samples were prepared for TEM imaging as described in (*62*) and (*63*). Briefly, samples were first fixed in a 2.5% glutaraldehyde solution (GA)(TAAB) in 0.1M phosphate buffer pH 7.4 (PB) at room temperature for an hour, then post fixation was done using 1,5 % osmium tetroxide (TAAB) and 1.5% potassium hexacyanoferrate (II)trihydrate (Sigma-Aldrich) in PB for an hour. The samples were then dehydrated by incubating them in a graded ethanol series (30%, 50%, 70%, 80%, 95%, and 100%) for 15 minutes at each concentration. Following dehydration, resin infiltration was done using Spurr resin (TAAB) with successive incubations in 33% and 66% resin for 2 hours each, and then 100% resin overnight. Finally, the resin was polymerized at 60°C overnight. Once resin polymerized, the sample blocks were trimmed and sectioned at an oblique angle using ultramicrotome (Leica EM FC) with a diamond knife. Then, the thin sections (70nm) on grid were first incubated with 1% uranyl acetate in darkness for 35 minutes and then incubated with lead citrate for 6 minutes.

All the TEM images were acquired using FEI Talos L 120C (accelerating voltage 120 kV). Velox data acquisition software was used with the following settings: image size 4096 pixels x 4096 pixels with different magnifications (2000X, 6700X, 11000X, 28000X) and 1s exposure time to acquire the images. The corresponding pixel size of each magnification were as follows: 6.929 nm at 2000X, 2.147 nm at 6700x, 1.345 nm at 11000X and 506.2 pm at 28000X.

### Cell Wall Thickness Quantification

Cell wall thickness measurements were done on TEM images taken on longitudinal cross sections across outer epidermal edges of transverse junctions on 4-days-old dark-grown Col-0 hypocotyls. Ten TEM images from five individual samples coming from 3 biological replicates were used. Briefly, using Fiji (*59*), the cell wall regions of the images were first manually segmented using the polygon selection tool followed by the application of the Fill and Clear Outside functions (*Edit>Fill* and *Edit>Clear Outside*) generating a binary image of the cell wall domain. The Local Thickness (*Analyze>Local Thickness>Local Thickness (complete process)*) function of Fiji was then applied to the binary images. The method is based on the computation of largest inscribed circle throughout the binary shape yielding a local thickness map. To quantify the local cell wall thickness in different subdomains of the segmented cell wall, we manually defined ROIs that were positioned on the Outer Epidermal Cell Wall (one for each side), Outer Epidermal Cell Edge and the Anticlinal Cell Wall and extracted local thickness values from the thickness map. Measurements were taken using the ROI manager in Fiji. Once all the measurements were completed for each cell wall domain, the average thickness for each cell wall subdomains and the aspect ratios between subdomains were calculated in Excel.

### Micro extensometer experiment

The extensometer used for the cell separation experiment was previously described in (*64*). The device consists of a 10g Futek load cell for force measurement, and two SmarAct nanopositioners to create movement in two axes. The software used to control the device was the MorphRobotX (MRX) framework of MorphoDynamX (morphographx.org/morphorobotx/). For this experiment, the micro extensometer was placed under a Nikon LV100ND Crest Optics Cicero spinning disk microscope and images were acquired as described in the corresponding confocal microscopy section.

The seedlings were mounted on the extensometer arms using PolySilTM S1001 silicone transfer adhesive and Tough Tags (Diversified Biotech) as described in (*64*) Using a pipette, a single drop of 0.2mg/ml propidium iodide (PI) (Sigma-Aldrich) prepared in Milli-Q water was placed on the hypocotyl (in the gap region between the tough tags). The PI solution was removed after 2.5 minutes, and the sample was immediately immersed in MiliQ Water.

After placing and staining the sample with PI as mentioned above, the first confocal image was acquired while the samples were held with 0mN of force. Then we exerted 10mN of force (stretching) on the hypocotyl using the *processes/Tools/MorphoRobotX/Experiment/Force based movement* process and acquired a Z-stack of the same region of interest. In the following steps, a 2mN force increment followed by confocal Z-stack acquisition, were performed until sample rupture. Note that between each force increment and confocal imaging, once the target force is reached, we waited 30 seconds before inactivating the extensometer arm and acquiring the Z-stack.

### Finite Element Modeling of hypocotyl epidermal cells

The geometries used for the finite element modeling (FEM) simulations were constructed using GMSH (v4.12.0; (*65*). Four different configurations were created to represent various mechanical scenarios in the hypocotyl epidermis, as described in details below.

All simulations were conducted using the BVPy modeling framework (*66*), a Python library built on FEniCS (*67*). The variational formulation employed a hyper-elastic model following the Saint-Venant Kirchhoff model.

In terms of material properties, the Poisson’s ratio for all subdomains was set to 0.3, and specific Young’s modulus are described for each configuration and scenario below.

Simulation outputs, including displacement fields, strain, and stress distributions, were saved in xdmf format for detailed post-processing. Visualization was performed using Paraview (v5.11.1).

- *Configuration 1: cross section through a group of fully adhering turgid cells (Fig 1 B)*

The geometry featured a cross-section of a group of turgid cells. The initial geometry was created using a hexagonal tiling approach, where hexagonal polygons with a radius of 3 µm were arranged to form a honeycomb pattern. A central cell and the following 4 surrounding layers of hexagonal cells around it were selected to be part of the study group of cells. To approximate a more natural cell arrangement, the outermost points of the mesh were snapped onto a circle with a 21 µm radius. To ensure a biologically relevant cell shape, the polygonal cells were then smoothed using the ksmooth method (smoothr 1.0.1) to reduce geometric sharp corner while preserving cell shape. Finally, all polygons were uniformly shrunk by 0.2 µm, resulting in a cell wall thickness of 0.4 µm. The cell wall domain was assigned a Young’s modulus of 0.05 MPa. Boundary conditions included Neumann boundary conditions applied to all inner edges of the mesh to create a normal force representing a turgor pressure of 0.5 MPa in all cells. A Dirichlet boundary condition imposing a fixed cell wall on the central cell was implemented to anchor the mesh and increase the simulation accuracy.

- *Configuration 2: epidermis surface mesh with staggered cell files and adhesive interfaces (Fig 1 C)*

The geometry featured staggered cell files segmented into two subdomains: the cells and the interface between cells.

Simulations explored two mechanical scenarios: i) Young’s modulus of the interface was set to half of the cell domain (5 MPa vs. 10 MPa); ii) Young’s modulus of the interface was doubled compared to the cell domain (10 MPa vs. 5 MPa) in order to simulate a doubled theoretical thickness. Boundary conditions applied a forced displacement at each end of the mesh using a Dirichlet boundary condition with displacements of 2.5 µm. resulting in a total strain of 3.2851e-2.

- *Configuration 3: longitudinal section mesh with two turgid cells and adhesive interfaces (Fig 1 D)*

A 2D mesh of a longitudinal section of the hypocotyl epidermis was created, consisting of two subdomains: the cell wall and the adhesive layer at the interface. Mechanical properties were assigned as follows: The cell wall subdomain was assigned a Young’s modulus of 10 MPa. The adhesive layer subdomain was assigned a Young’s modulus of 5 MPa. Boundary conditions included a Dirichlet boundary conditions imposing displacements of 0.5 µm at both ends of the mesh, resulting in a total strain of 0.025. In addition, Neumann boundary conditions was applied on all inner edges of the mesh to model cell turgor pressure of 0.5 MPa.

- *Configuration 4: longitudinal section mesh with two turgid cells and four subdomains outer epidermal edge (Fig 4 A)*

A more detailed 2D mesh was created, representing a longitudinal section of the hypocotyl epidermis. This geometry included four subdomains at the outer epidermal edge: the supracellular outer epidermal wall (SOEW), the outer epidermal edge filling (OEEF), the inner walls (IW), and the middle lamella (ML). The relative thicknesses of these subdomains were informed by measurements from a TEM longitudinal section image (Fig 3 H) (*59*).

Simulations explored five mechanical scenarios: i) uniform mechanical properties across all subdomains with a Young’s modulus of 10 MPa; ii) a softer middle lamella with a Young’s modulus of 1 MPa compared to 10 MPa for other subdomains; iii) a softer outer epidermal edge filling with a Young’s modulus of 1 MPa; iv) a softer supracellular outer epidermal wall with a Young’s modulus of 1 MPa; v) both the supracellular outer epidermal wall and the outer epidermal edge filling with a Young’s modulus of 1 MPa, while other subdomains were set to 10 MPa.

Boundary conditions included a Dirichlet boundary condition imposing displacements of 0.25 µm at each end of the mesh, resulting in a total strain of 6.3532e-3., In addition, Neumann boundary conditions of 0.2 MPa was applied on all inner edges of the mesh to model cell turgor pressure.

## References

1. T. J. C. Harris, U. Tepass, Adherens junctions: from molecules to morphogenesis. Nat Rev Mol Cell Biol 11, 502–514 (2010).

2. E. Zamir, B. Geiger, J. E. Schaffer, “Cell-Cell Interaction | Focal Adhesions and Related Integrin Contacts” in Encyclopedia of Biological Chemistry III (Elsevier, 2021; https://linkinghub.elsevier.com/retrieve/pii/B9780128194607001584), pp. 716–721.

3. A. Atakhani, L. Bogdziewiez, S. Verger, Characterising the mechanics of cell–cell adhesion in plants. Quant Plant Bio. 3, e2 (2022).

4. A. I. Baba, S. Verger, Cell adhesion maintenance and controlled separation in plants. *Front*. Plant Physiol. 2, 1369575 (2024).

5. F. B. Daher, S. A. Braybrook, How to let go: pectin and plant cell adhesion. Front. Plant Sci. 6 (2015).

6. M. C. Jarvis, S. P. H. Briggs, J. P. Knox, Intercellular adhesion and cell separation in plants. Plant Cell & Environment 26, 977–989 (2003).

7. J. P. Knox, Cell adhesion, cell separation and plant morphogenesis. The Plant Journal 2, 137–141 (1992).

8. C. T. Anderson, J. J. Kieber, Dynamic Construction, Perception, and Remodeling of Plant Cell Walls. Annu. Rev. Plant Biol. 71, 39–69 (2020).

9. N. Hoffmann, S. King, A. L. Samuels, H. E. McFarlane, Subcellular coordination of plant cell wall synthesis. Developmental Cell 56, 933–948 (2021).

10. A. R. Paredez, C. R. Somerville, D. W. Ehrhardt, Visualization of Cellulose Synthase Demonstrates Functional Association with Microtubules. Science 312, 1491–1495 (2006).

11. D. J. Cosgrove, Structure and growth of plant cell walls. Nat Rev Mol Cell Biol 25, 340–358 (2024).

12. T. Zhang, Y. Zheng, D. J. Cosgrove, Spatial organization of cellulose microfibrils and matrix polysaccharides in primary plant cell walls as imaged by multichannel atomic force microscopy. The Plant Journal 85, 179–192 (2016).

13. A. J. Bidhendi, O. Lampron, F. P. Gosselin, A. Geitmann, Cell geometry regulates tissue fracture. Nat Commun 14, 8275 (2023).

14. R. B. Malek, G. Mouille, “The Middle Lamella between Anthropic and Evolutionary Contexts” in Plant Cell Walls (2023), p. 10.

15. M. S. Zamil, A. Geitmann, The middle lamella—more than a glue. Phys. Biol. 14, 015004 (2017).

16. S. Verger, S. Chabout, E. Gineau, G. Mouille, Cell adhesion in plants is under the control of putative O-fucosyltransferases. Development, dev.132308 (2016).

17. J. Du, A. Kirui, S. Huang, L. Wang, W. J. Barnes, S. N. Kiemle, Y. Zheng, Y. Rui, M. Ruan, S. Qi, S. H. Kim, T. Wang, D. J. Cosgrove, C. T. Anderson, C. Xiao, Mutations in the Pectin Methyltransferase QUASIMODO2 Influence Cellulose Biosynthesis and Wall Integrity in Arabidopsis. The Plant Cell 32, 3576–3597 (2020).

18. R. Lorrai, F. Francocci, K. Gully, H. J. Martens, G. De Lorenzo, C. Nawrath, S. Ferrari, Impaired Cuticle Functionality and Robust Resistance to Botrytis cinerea in Arabidopsis thaliana Plants With Altered Homogalacturonan Integrity Are Dependent on the Class III Peroxidase AtPRX71. Front. Plant Sci. 12, 696955 (2021).

19. H.-T. Cho, D. J. Cosgrove, Altered expression of expansin modulates leaf growth and pedicel abscission in *Arabidopsis thaliana*. Proc. Natl. Acad. Sci. U.S.A. 97, 9783–9788 (2000).

20. S. Y. Rhee, E. Osborne, P. D. Poindexter, C. R. Somerville, Microspore Separation in the *quartet 3* Mutants of Arabidopsis Is Impaired by a Defect in a Developmentally Regulated Polygalacturonase Required for Pollen Mother Cell Wall Degradation. Plant Physiology 133, 1170–1180 (2003).

21. S. Cai, C. C. Lashbrook, Stamen Abscission Zone Transcriptome Profiling Reveals New Candidates for Abscission Control: Enhanced Retention of Floral Organs in Transgenic Plants Overexpressing Arabidopsis *ZINC FINGER PROTEIN2*. Plant Physiology 146, 1305–1321 (2008).

22. G. Drakakaki, Polysaccharide deposition during cytokinesis: Challenges and future perspectives. Plant Science 236, 177–184 (2015).

23. F. Miart, T. Desprez, E. Biot, H. Morin, K. Belcram, H. Höfte, M. Gonneau, S. Vernhettes, Spatio-temporal analysis of cellulose synthesis during cell plate formation in A rabidopsis. The Plant Journal 77, 71–84 (2014).

24. C. Van Oostende-Triplet, D. Guillet, T. Triplet, E. Pandzic, P. W. Wiseman, A. Geitmann, Vesicle Dynamics during Plant Cell Cytokinesis Reveals Distinct Developmental Phases. Plant Physiol. 174, 1544–1558 (2017).

25. S. A. Braybrook, H. Jönsson, Shifting foundations: the mechanical cell wall and development. Current Opinion in Plant Biology 29, 115–120 (2016).

26. D. Kierzkowski, A.-L. Routier-Kierzkowska, Cellular basis of growth in plants: geometry matters. Current Opinion in Plant Biology 47, 56–63 (2019).

27. C. Kirchhelle, C.-M. Chow, C. Foucart, H. Neto, Y.-D. Stierhof, M. Kalde, C. Walton, M. Fricker, R. S. Smith, A. Jérusalem, N. Irani, I. Moore, The Specification of Geometric Edges by a Plant Rab GTPase Is an Essential Cell-Patterning Principle During Organogenesis in Arabidopsis. Developmental Cell 36, 386–400 (2016).

28. M. Majda, N. Trozzi, G. Mosca, R. S. Smith, How Cell Geometry and Cellular Patterning Influence Tissue Stiffness. IJMS 23, 5651 (2022).

29. M. Asaoka, M. Ooe, S. Gunji, P. Milani, G. Runel, G. Horiguchi, O. Hamant, S. Sawa, H. Tsukaya, A. Ferjani, Stem integrity in *Arabidopsis thaliana* requires a load-bearing epidermis. Development 148, dev198028 (2021).

30. R. Kelly-Bellow, K. Lee, R. Kennaway, J. E. Barclay, A. Whibley, C. Bushell, J. Spooner, M. Yu, P. Brett, B. Kular, S. Cheng, J. Chu, T. Xu, B. Lane, J. Fitzsimons, Y. Xue, R. S. Smith, C. D. Whitewoods, E. Coen, Brassinosteroid coordinates cell layer interactions in plants via cell wall and tissue mechanics. Science 380, 1275–1281 (2023).

31. U. Kutschera, The Growing Outer Epidermal Wall: Design and Physiological Role of a Composite Structure. Annals of Botany 101, 615–621 (2008).

32. T. I. Baskin, O. E. Jensen, On the role of stress anisotropy in the growth of stems. Journal of Experimental Botany 64, 4697–4707 (2013).

33. S. Robinson, C. Kuhlemeier, Global Compression Reorients Cortical Microtubules in Arabidopsis Hypocotyl Epidermis and Promotes Growth. Current Biology 28, 1794–1802.e2 (2018).

34. S. Verger, Y. Long, A. Boudaoud, O. Hamant, A tension-adhesion feedback loop in plant epidermis. eLife 7, e34460 (2018).

35. J. Mathur, N. Mathur, B. Kernebeck, M. Hülskamp, Mutations in Actin-Related Proteins 2 and 3 Affect Cell Shape Development in Arabidopsis. Plant Cell 15, 1632–1645 (2003).

36. W. G. T. Willats, C. Orfila, G. Limberg, H. C. Buchholt, G.-J. W. M. Van Alebeek, A. G. . Voragen, S. E. Marcus, T. M. I. E. Christensen, J. D. Mikkelsen, B. S. Murray, J. P. Knox, Modulation of the Degree and Pattern of Methyl-esterification of Pectic Homogalacturonan in Plant Cell Walls. Journal of Biological Chemistry 276, 19404–19413 (2001).

37. R. Galletti, S. Verger, O. Hamant, G. C. Ingram, Developing a ‘thick skin’: a paradoxical role for mechanical tension in maintaining epidermal integrity? Development 143, 3249–3258 (2016).

38. C. D. Whitewoods, Riddled with holes: Understanding air space formation in plant leaves. PLoS Biol 19, e3001475 (2021).

39. M. C. Jarvis, Intercellular separation forces generated by intracellular pressure. Plant Cell & Environment 21, 1307–1310 (1998).

40. J. P. Knox, Paul J. Linstead, J. King, C. Cooper, K. Roberts, Pectin esterification is spatially regulated both within cell walls and between developing tissues of root apices. Planta 181 (1990).

41. C. C. Parker, M. L. Parker, A. C. Smith, K. W. Waldron, Pectin Distribution at the Surface of Potato Parenchyma Cells in Relation to Cell−Cell Adhesion. J. Agric. Food Chem. 49, 4364–4371 (2001).

42. A. M. Catesson, J. C. Roland, Sequential Changes Associated with Cell Wall Formation and Fusion in the Vascular Cambium. IAWA J 2, 151–162 (1981).

43. C. E. Jeffree, J. E. Dale, S. C. Fry, The genesis of intercellular spaces in developing leaves ofPhaseolus vulgaris L. Protoplasma 132, 90–98 (1986).

44. D. Matar, A. M. Catesson, Cell plate development and delayed formation of the pectic middle lamella in root meristems. Protoplasma 146, 10–17 (1988).

45. L. Elliott, M. Kalde, A.-K. Schürholz, X. Zhang, S. Wolf, I. Moore, C. Kirchhelle, A self-regulatory cell-wall-sensing module at cell edges controls plant growth. Nat. Plants 10, 483–493 (2024).

46. L. Elliott, C. Kirchhelle, The importance of being edgy: cell geometric edges as an emerging polar domain in plant cells. Journal of Microscopy 278, 123–131 (2020).

47. A. Fruleux, S. Verger, A. Boudaoud, Feeling Stressed or Strained? A Biophysical Model for Cell Wall Mechanosensing in Plants. Front. Plant Sci. 10, 757 (2019).

48. R. Galletti, K. L. Johnson, S. Scofield, R. San-Bento, A. M. Watt, J. A. H. Murray, G. C. Ingram, DEFECTIVE KERNEL 1 promotes and maintains plant epidermal differentiation. Development 142, 1978–1983 (2015).

49. C. Kirchhelle, O. Hamant, Discretizing the cellular bases of plant morphogenesis: Emerging properties from subcellular and noisy patterning. Current Opinion in Cell Biology 81, 102159 (2023).

50. D. Tran, R. Galletti, E. D. Neumann, A. Dubois, R. Sharif-Naeini, A. Geitmann, J.-M. Frachisse, O. Hamant, G. C. Ingram, A mechanosensitive Ca2+ channel activity is dependent on the developmental regulator DEK1. Nat Commun 8, 1009 (2017).

51. N. Hoffmann, E. Mohammad, H. E. McFarlane, Disrupting cell wall integrity impacts endomembrane trafficking to promote secretion over endocytic trafficking. Journal of Experimental Botany 75, 3731–3747 (2024).

52. E. C. Cocking, Some effects on roots of tomato seedlings. Biochemistry Journal 76, 51–52 (1960).

53. G. Mouille, M. Ralet, C. Cavelier, C. Eland, D. Effroy, K. Hématy, L. McCartney, H. N. Truong, V. Gaudon, J. Thibault, A. Marchant, H. Höfte, Homogalacturonan synthesis in *Arabidopsis thaliana* requires a Golgi-localized protein with a putative methyltransferase domain. The Plant Journal 50, 605–614 (2007).

54. J. Le, S. E.-D. El-Assal, D. Basu, M. E. Saad, D. B. Szymanski, Requirements for Arabidopsis ATARP2 and ATARP3 during Epidermal Development. Current Biology 13, 1341–1347 (2003).

55. S. Li, L. Blanchoin, Z. Yang, E. M. Lord, The Putative Arabidopsis Arp2/3 Complex Controls Leaf Cell Morphogenesis. Plant Physiology 132, 2034–2044 (2003).

56. Ö. Erguvan, M. Louveaux, O. Hamant, S. Verger, ImageJ SurfCut: a user-friendly pipeline for high-throughput extraction of cell contours from 3D image stacks. BMC Biol 17, 38 (2019).

57. A. J. Bidhendi, Y. Chebli, A. Geitmann, Fluorescence visualization of cellulose and pectin in the primary plant cell wall. Journal of Microscopy 278, 164–181 (2020).

58. E. H. W. Meijering, W. J. Niessen, M. A. Viergever, Quantitative evaluation of convolution-based methods for medical image interpolation. Medical Image Analysis 5, 111–126 (2001).

59. J. Schindelin, I. Arganda-Carreras, E. Frise, V. Kaynig, M. Longair, T. Pietzsch, S. Preibisch, C. Rueden, S. Saalfeld, B. Schmid, J.-Y. Tinevez, D. J. White, V. Hartenstein, K. Eliceiri, P. Tomancak, A. Cardona, Fiji: an open-source platform for biological-image analysis. Nat Methods 9, 676–682 (2012).

60. J. E. Sader, R. Borgani, C. T. Gibson, D. B. Haviland, M. J. Higgins, J. I. Kilpatrick, J. Lu, P. Mulvaney, C. J. Shearer, A. D. Slattery, P.-A. Thorén, J. Tran, H. Zhang, H. Zhang, T. Zheng, A virtual instrument to standardise the calibration of atomic force microscope cantilevers. Review of Scientific Instruments 87, 093711 (2016).

61. M. Seale, A. Kiss, S. Bovio, I. M. Viola, E. Mastropaolo, A. Boudaoud, N. Nakayama, Dandelion pappus morphing is actuated by radially patterned material swelling. Nat Commun 13, 2498 (2022).

62. M. Barberon, J. E. M. Vermeer, D. De Bellis, P. Wang, S. Naseer, T. G. Andersen, B. M. Humbel, C. Nawrath, J. Takano, D. E. Salt, N. Geldner, Adaptation of Root Function by Nutrient-Induced Plasticity of Endodermal Differentiation. Cell 164, 447–459 (2016).

63. A. Berhin, D. De Bellis, R. B. Franke, R. A. Buono, M. K. Nowack, C. Nawrath, The Root Cap Cuticle: A Cell Wall Structure for Seedling Establishment and Lateral Root Formation. Cell 176, 1367–1378.e8 (2019).

64. A. I. Baba, L. Lisica, A. Atakhani, B. Aryal, L. Bogdziewiez, Ö. Erguvan, A. Heymans, P. K. Jewaria, R. S. Smith, R. P. Bhalerao, S. Verger, Rhamnogalacturonan-II dimerization deficiency impairs the coordination between growth and adhesion maintenance in plants. Plant Biology [Preprint] (2024). 10.1101/2024.11.26.625362.

65. C. Geuzaine, J. Remacle, Gmsh: A 3-D finite element mesh generator with built-in pre- and post-processing facilities. Numerical Meth Engineering 79, 1309–1331 (2009).

66. F. Gacon, C. Godin, O. Ali, BVPy: A FEniCS-based Python package to ease the expression and study of boundary value problems in Biology. JOSS 6, 2831 (2021).

67. A. Logg, K.-A. Mardal, G. Wells, Eds., Automated Solution of DiSerential Equations by the Finite Element Method: The FEniCS Book (Springer Berlin Heidelberg, Berlin, Heidelberg, 2012; https://link.springer.com/10.1007/978-3-642-23099-8)vol. 84 of Lecture Notes in Computational Science and Engineering.

