## Supplemental figures S1-S3 and table S1 for "Outer epidermal edges mediate cell-cell adhesion for tissue integrity in plants"

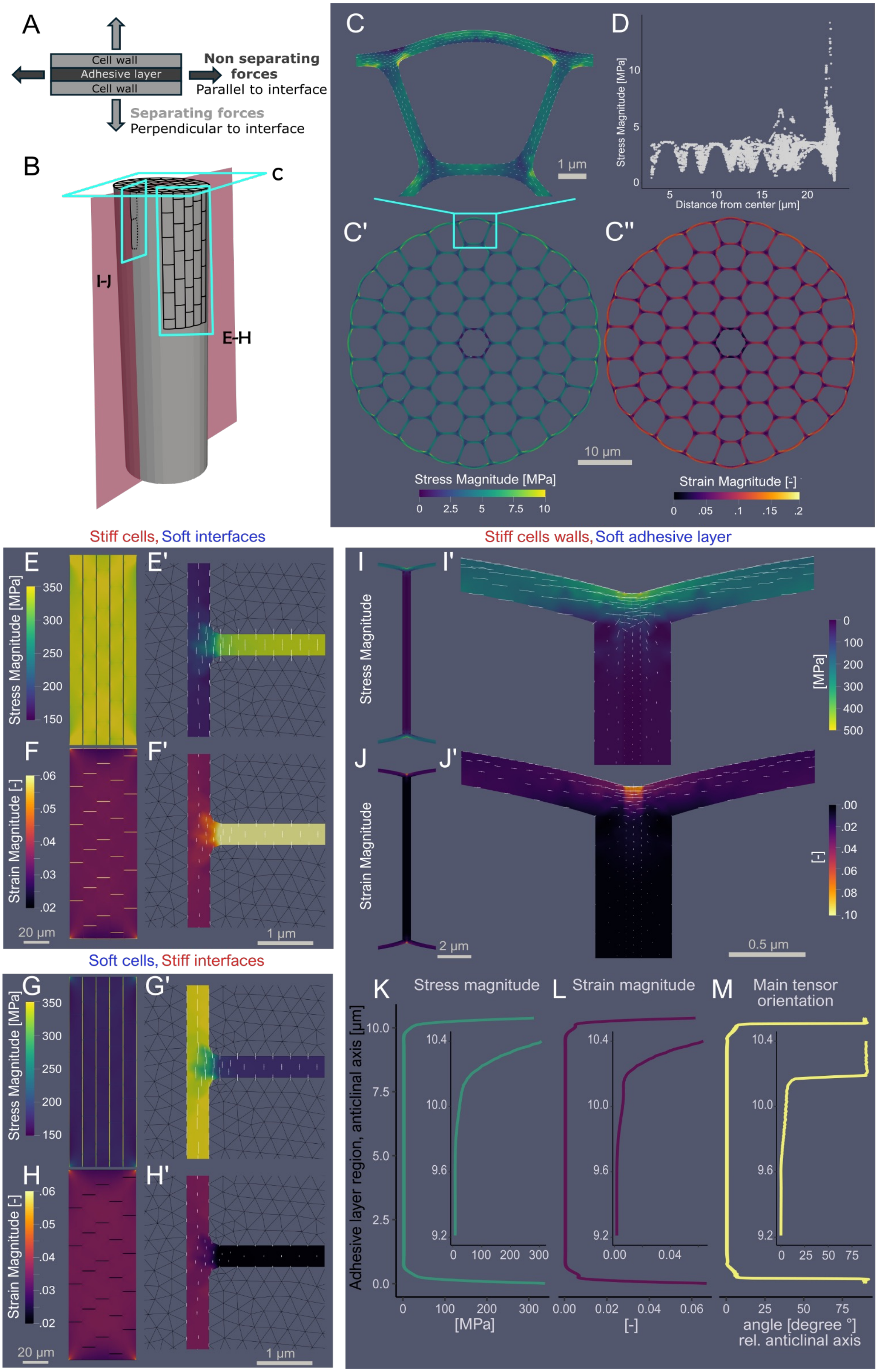

A
Cell wall
Adhesive layer
Cell wall
Non separating forces
Parallel to interface
Separating forces
Perpendicular to interface
B
C
I-J
E-H
C
D
1 µm
Stress Magnitude [MPa]
Distance from center [µm]
C'
C''
10 µm
Stress Magnitude [MPa]
0 2.5 5 7.5 10
Strain Magnitude [-]
0 .05 .1 .15 .2
Stiff cells, Soft interfaces
E
E'
Stress Magnitude [MPa]
F
F'
Strain Magnitude [-]
20 µm
1 µm
Stiff cells walls, Soft adhesive layer
I
I'
Stress Magnitude
[MPa]
J
J'
Strain Magnitude
[-]
2 µm
0.5 µm
Soft cells, Stiff interfaces
G
G'
Stress Magnitude [MPa]
H
H'
Strain Magnitude [-]
20 µm
1 µm
K
Stress magnitude
L
Strain magnitude
M
Main tensor orientation
Adhesive layer region, anticlinal axis [µm]
[MPa]
[-]
angle [degree °] rel. anticlinal axis

Figure S1: Additional simulation output supporting figure 1. (A) Conceptual representation of the "adhesive layer model." (B) Conceptual schematic illustrating different planes and viewpoints used to analyse a cylindrical plant organ. (C) Magnified view of the stress heatmap in an epidermal cell from a 2D mesh representing a cross-section through a group of turgid cells. The white segments indicating major and minor tensor orientations. (C’) Global view after displacement of the 2D mesh cross-section with the stress heatmap. (C”) Global view after displacement of the same 2D mesh cross-section with a strain heatmap. (D) Scatter plot of stress magnitude as a function of the distance from the center of the 2D cross-section mesh. (E) Stress heatmap of the epidermal surface mesh under tensile stretching, where the Young's modulus of the interfaces was set to half of the cell domains. (F) Strain heatmap corresponding to (E) under the same conditions. (E’ & F’) Close-up view of the interface at a tri-cellular junction, with white segments marking principal tensor orientations. (G) Stress heatmap of the epidermal surface mesh under tensile stretching, where the Young’s modulus of the cell domain was set to half of the interface. (H) Strain heatmap corresponding to (G) under the same conditions. (G’ & H’) Close-up view of the interface at a tri-cellular junction, with white segments marking principal tensor orientations. Note that in (E’-H’) for illustration purposes, we only display stress distribution in the adhesive layer. (I) Stress heatmap on an anticlinal wall of the 2D mesh representing the longitudinal section of epidermal cells under tensile stretch and turgor pressure, where the Young’s modulus of the adhesive layer was set to half of the cell wall domains. (I’) Close-up stress heatmap of the outer epidermal edge, with white segments marking principal tensor orientations. (J) Strain heatmap corresponding to (I) under the same conditions. (J’) Close-up strain heatmap of the outer epidermal edge, with white segments marking principal tensor orientations. (K) Stress magnitude [MPa] along the anticlinal axis within the adhesive layer subdomain. (L) Strain magnitude [-] along the anticlinal axis within the adhesive layer subdomain. (M) Principal tensor orientation [deg. °; relative to the anticlinal axis ] within the adhesive layer subdomain.

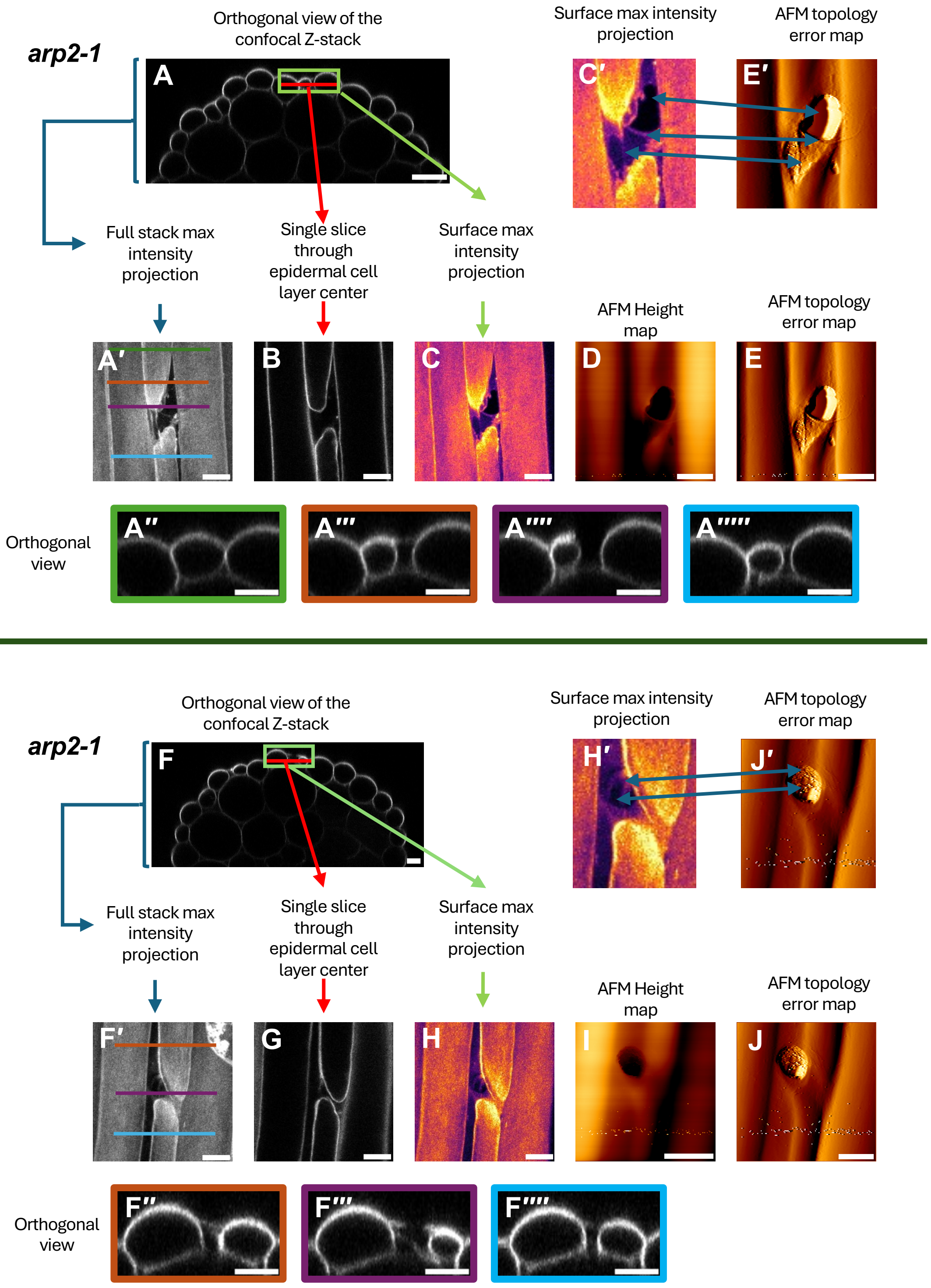

arp2-1
Orthogonal view of the confocal Z-stack
A
Surface max intensity projection
AFM topology error map
C′
E′
Full stack max intensity projection
Single slice through epidermal cell layer center
Surface max intensity projection
AFM Height map
AFM topology error map
A′
B
C
D
E
Orthogonal view
A″
A‴
A⁗
A″‴
arp2-1
Orthogonal view of the confocal Z-stack
F
Surface max intensity projection
AFM topology error map
H′
J′
Full stack max intensity projection
Single slice through epidermal cell layer center
Surface max intensity projection
AFM Height map
AFM topology error map
F′
G
H
I
J
Orthogonal view
F″
F‴
F⁗

Figure S2: Correlative confocal-AFM confirms partial material continuity across mildly separated cells. Two examples highlighting outer epidermal wall continuity across mildly separated cells in arp2-1. (A and F) Representative orthogonal view of the confocal Z-stack images from 4-day-old dark-grown *arp2-1* hypocotyls stained with propidium iodide. Red lines in A and F show where the single slice images generated for B and H respectively. (A′ and F′) Representative full stack max intensity projections confocal images. (A″- A″″ and F″-F″″) Representative Single slice orthogonal images where the frame colors indicates where the orthogonal images where sampled from in A′ and F′ (note colored lines in A′ and F′). (B and G) Representative single slice confocal images throough the epidermal layer at the mildly separated junction. Red lines in A and F indicate where the single slice images coming from. (C and H) Representative surface max intensity projection confocal images from a sub portion of the stack at the surface (green rectangles in A and F show the cropped regions and the depth of the images). Note: we applied a "fire" Look up table to highlight the similarities with the AFM topology error map images. (C′ and H′) cropped from C and H. (D-E and I-J) AFM images from the same regions as imaged with confocal microscopy. (D and I) Height maps. (E and J) Topology (error maps) images. (E′ and J′) cropped from E and J. Arrows between C'/E' and H'/J', highlight the similiarity of the signal detected with PI staining and the AFM surface topology. Scale bars: A-C 20µm, A′-G′, A″-A″″ and G″-G″″ 10µm, E and K 5µm.

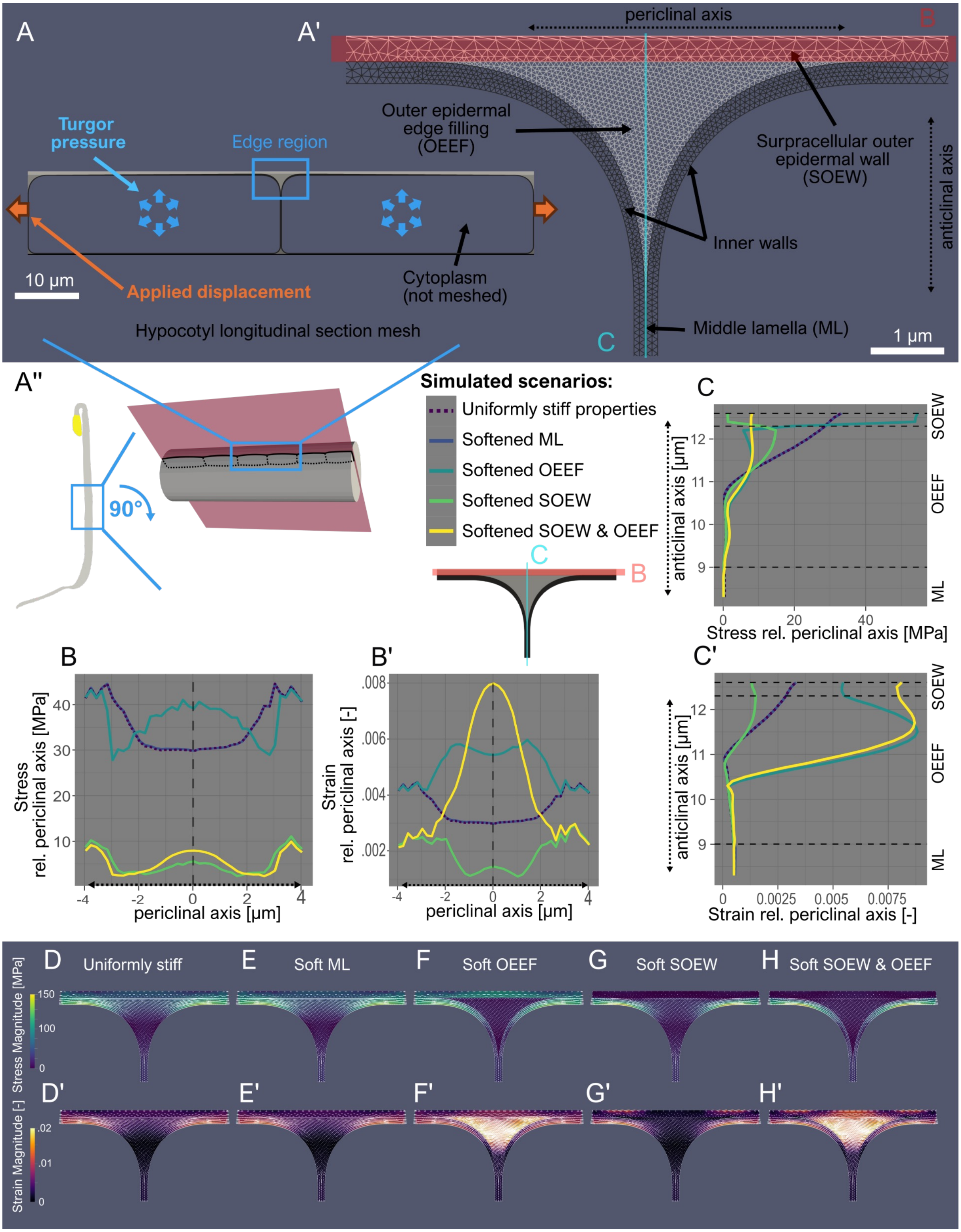
A
A'
periclinal axis
B
Outer epidermal edge filling (OEEF)
Surpracellular outer epidermal wall (SOEW)
anticlinal axis
Turgor pressure
Edge region
Inner walls
Cytoplasm (not meshed)
10 µm
Applied displacement
Hypocotyl longitudinal section mesh
C
Middle lamella (ML)
1 µm
A''
90°
Simulated scenarios:
Uniformly stiff properties
Softened ML
Softened OEEF
Softened SOEW
Softened SOEW & OEEF
Stress rel. periclinal axis [MPa]
anticlinal axis [µm]
SOEW
OEEF
ML
Stress rel. periclinal axis [MPa]
periclinal axis [µm]
B'
Strain rel. periclinal axis [-]
C'
Strain rel. periclinal axis [-]
D Uniformly stiff
E Soft ML
F Soft OEEF
G Soft SOEW
H Soft SOEW & OEEF
Stress Magnitude [MPa]
Strain Magnitude [-]
D'
E'
F'
G'
H'

Figure S3: Additional output and quantifications supporting figure 4. Finite Element Method (FEM) analysis of stress and strain distribution on a longitudinal section of hypocotyl epidermal cells under varied subdomain Young's modulus conditions. (A) 2D mesh of the hypocotyl longitudinal section, with (A') a close-up of the edge region at the cell junction. Four subdomains are identified: the supracellular outer epidermal wall (SOEW), the outer epidermal edge filling (OEEF), the inner walls, and the middle lamella (ML). (A") Conceptual schematic illustrating planes and viewpoints used to analyse a dark-grown hypocotyl epidermal cell under tensile stretch and turgor pressure. Five simulated scenarios were designed with varied Young's modulus values. Unless otherwise specified, Young's modulus for all subdomains was set to 10 MPa, with specific scenarios softening subdomains to 1 MPa. (B-C) To better understand the forces that would threaten to separate cells, we generated graphs where we extracted the component of the stress and strain that is parallel to the stretching applied. (B) Stress magnitude [MPa] and (B') strain magnitude [-] along the periclinal axis within the SOEW subdomain (Red region on the mesh in A'). (C) Stress magnitude [MPa] and (C') strain magnitude [-] along the anticlinal axis within the cell junction (Cyan region on the mesh in A'). The cell junction area encompasses nodes within 0.1625 µm of the median line separating the two cells (inner walls and ML).(D–H) Close-ups of the edge region, showing stress distribution with tensor orientations under different conditions:(D) Uniform properties; (E) Softened ML; (F) Softened OEEF; (G) Softened SOEW; (H) Softened OEEF and SOEW. (D'–H') Corresponding strain magnitude heatmaps for each scenario.

Table S1: Cell wall thickness measurements in 4-day-old dark-grown hypocotyl of wild type (Col-0). OECW: Outer Epidermal cell wall; CW: Cell Wall; OEE: Outer epidermal Edges. See corresponding representations in figure 3. Values of Right OECW and Left OECW were averaged to yield the "Average OECW" value. This data set is plotted in Figure 3 H.

| Sample | Right OECW (µm) | Left OECW (µm) | Anticlinal CW (µm) | OEE (µm) | Average OECW (µm) | Ratio OECW/ Anticlinal CW | Ratio OEE/ OECW | Ratio OEE/ Anticlinal CW |
|---|---|---|---|---|---|---|---|---|
| Col-0_R1_S2_J1 | 0,355 | 0,366 | 0,198 | 1,019 | 0,361 | 1,822 | 2,826 | 5,150 |
| Col-0_R2_S9522_J1 | 0,946 | 0,943 | 0,336 | 1,968 | 0,944 | 2,810 | 2,975 | 5,855 |
| Col-0_R2_S9522_J2 | 0,664 | 0,834 | 0,311 | 1,886 | 0,749 | 2,411 | 2,518 | 6,070 |
| Col-0_R3_S9595_J1 | 0,809 | 0,964 | 0,324 | 1,768 | 0,887 | 2,732 | 1,995 | 5,450 |
| Col-0_R3_S9595_J2 | 0,388 | 0,328 | 0,180 | 0,932 | 0,358 | 1,986 | 2,602 | 5,168 |
| Col-0_R3_S9690_J1 | 0,392 | 0,411 | 0,210 | 1,134 | 0,401 | 1,916 | 2,825 | 5,412 |
| Col-0_R3_S9690_J2 | 0,589 | 0,541 | 0,142 | 1,481 | 0,565 | 3,966 | 2,622 | 10,400 |
| Col-0_R3_S9690_J3 | 0,384 | 0,409 | 0,242 | 1,248 | 0,396 | 1,641 | 3,149 | 5,169 |
| Col-0_R3_S9690_J5 | 0,729 | 0,635 | 0,220 | 1,324 | 0,682 | 3,101 | 1,942 | 6,022 |
| Col-0_R3_S9694_J1 | 0,439 | 0,451 | 0,205 | 1,172 | 0,445 | 2,166 | 2,636 | 5,708 |
| **Average** | | | **0,237** | **1,393** | **0,579** | **2,455** | **2,609** | **6,040** |
